# Effects of altered gravity on adrenergic-mediated cAMP signalling in intact cells

**DOI:** 10.1101/2025.03.02.640942

**Authors:** Marc Bathe-Peters, Iqra Sohail, Alexei Sirbu, Katharina Schneider, Tommaso Patriarchi, Anantha Anilkumar, Yannick Lichterfeld, Christian Liemersdorf, Primal de Lanerolle, Paolo Annibale

**Author notes:** equally contributing authors (Marc Bathe-Peters, Iqra Sohail).

## Abstract

Spaceflight-induced cardiac atrophy and rhythm disorders are linked to dysregulation of the adrenergic-cAMP-PKA pathway. Gravity-dependent alterations in adrenergic signaling, particularly cAMP dynamics, remain poorly understood. Using fluorescence biosensors, we studied intact cells under simulated microgravity and hypergravity. We observed shifts in the EC50 of cAMP production: leftward under hypergravity and rightward in microgravity, with faster cAMP accumulation kinetics in hypergravity. Cytoskeletal remodelling, hypothesized to be a determinant of such chances, was negligible, suggesting alternative mechanisms. These findings highlight significant gravity-induced offsets in the pharmacology of a prototypical GPCR, with implications not only for adrenergic signalling but also for other pathways of pharmacological interest, potentially informing countermeasures for astronaut health and pharmacology in altered gravity settings.

## Introduction

Muscle atrophy, including cardiac atrophy, sporadically accompanied by cardiac rhythm disorders, is a common side effect after prolonged spaceflight. Moreover, orthostatic intolerance and overall cardiac deconditioning is observed upon re-entry to 1g conditions (Auber 2005; Jung et al. 2005; Goswami et al. 2021). The adrenergic-cAMP-Protein kinase A (PKA) pathway, in regulating inotropic response of the heart to catecholamines, has a deep-rooted role in modulating these processes (Lefkowitz, Rockman, and Koch 2000).

A large portion of extracellular signals in human cells, including catecholamines, are mediated by G protein-coupled receptors (GPCRs), a family of over 800 membrane proteins and a prominent drug target (Hilger, Masureel, and Kobilka 2018). Their function is regulated by a complex network of extra and intra-cellular interactions that is a prime candidate to be affected by the pleiotropic effects originating from altered gravity.

Alterations to GPCR-mediated signaling cascades by biophysical factors can be observed at the level of second messenger production, e.g. cAMP for the canonical adrenergic stimulatory pathway (Sirbu et al. 2024). Changes in intracellular cAMP have broad consequences on cellular homeostasis, often cell-type specific, such as PKA-mediated increase in heart rate, immune regulation, chromatin condensation and regulation of intracellular transport (Musheshe, Schmidt, and Zaccolo 2018).

Early reports have shown that cAMP homeostasis is affected in cellular organisms and single cells exposed to periods of altered gravity (Hemmersbach et al. 2002; Albi et al. 2011), with results in single cells showing a decrease in cAMP-mediated response in microgravity. Nevertheless, further to pioneering late 1980s and early 1990s investigations addressing downstream PKA, the molecular mechanisms affected by altered gravity along this signaling axis remain rather unclear (Mednieks et al. 1987; Philpott 1990). More recent evidence based on RNA sequencing in non cardiac tissue, namely hematopoietic stem and progenitor cells that were exposed to microgravity in low earth orbit, indicated a depression of the cAMP-CREB axis (Wang et al. 2019).

However, thanks to optical technologies, most prominently fluorescence biosensors, it is now possible to directly visualize the activity of the molecular players in this pathway and explore how their function is affected by altered gravity conditions (Bock et al. 2020; Sirbu et al. 2024). This shall allow identifying and potentially to counteract at least some of the molecular determinants of cardiac deconditioning occurring in astronauts.

We conducted measurements in intact cells in simulated microgravity and hypergravity conditions, both in cells fixed immediately after altered gravity exposure, as well as validating some of these measurements by conducting live cell imaging readouts in hypergravity. Our data indicate a shift of the EC_50_ of cAMP production to the left for increasing periods of 2g exposure, and a shift to the right in microgravity conditions. Moreover, real-time kinetics of cAMP accumulation in cells upon catecholamine stimulation are also faster in cells exposed to hypergravity. Finally, we tested the hypothesis that the observed signaling behavior could be connected to remodeling of the cortical cytoskeleton (Gruener 1994; Crawford-Young 2006; Rosner et al. 2006; Wu et al. 2022), observing remarkably limited changes in cytoskeletal remodeling under the altered gravity conditions used in our experiments.

These results suggest that altered gravity induces an overall offset of GPCR-mediated pharmacology, reflected in alterations of cell-wide cAMP levels, and altered receptor activation kinetics. These results, validated in the important adrenergic signaling axis, have the potential to be extended to other important classes of receptors, mediating nociception, immune response, cell motility, to name only a few, with far reaching implications for pharmacology as well as -more broadly-for physiology in altered gravity settings.

## Results

We set out to measure the effect of altered gravity on the β-AR signaling cascade, by monitoring downstream cAMP production in HEK293 cells transiently expressing the FRET biosensor EPAC-S^H187^ (Klarenbeek et al. 2015), and monitoring FRET changes using a sensitized emission approach. Altered gravity conditions were achieved by either placing cells in a clinostat machine, which simulates microgravity (µg), or in a centrifuge that generates hypergravity conditions corresponding to 2g. In both instances we conducted incubations of 24h, while cells were maintained at 37°C and 5% CO_2_ conditions in incubators enclosing respectively the clinostats and the centrifuges.

For the cells exposed to microgravity conditions we conducted a concentration-response curve perfusing cells immediately after the simulated microgravity cycle with increasing concentrations of the β-AR agonist isoproterenol (**Figure 1a**). After five minutes incubation with the agonist, which typically corresponds to peak cAMP response (Sirbu et al. 2024), cells were fixed prior to subsequent imaging. Comparing the concentration-response curves for isoproterenol stimulations in live and fixed cells we showed that fixation using 4% paraformaldehyde and 0.2% glutaraldehyde does not affect the response of the biosensor as compared to the same readout conducted in living cells (**Figure S1a**). By imaging individual cells in an epifluorescence microscope equipped for sensitized emission FRET we could then map cAMP levels by means of the donor to FRET ratio, which changes in concert with intracellular cAMP levels. Our data show a right shift of the concentration-response curve, as the EC_50_ value (half maximal response to isoproterenol) goes from 10.6 nM of the ground control to 78.5 nM in microgravity (**Figure 1a**), as well as a reduction in maximal response (Emax). This observation means that in microgravity more drug is required to achieve an equivalent response to the control experiment, suggesting a blunting of the adrenergic-cAMP axis. We next moved on to explore the effects of hypergravity exposure on β-AR-mediated cAMP production. Cells transiently expressing the FRET EPAC-S^H187^ biosensor plated in 96-well microplates were now exposed to 24h of 2g hypergravity on the MuSIC centrifuge, stimulated for 5 minutes using isoproterenol, and then subsequently fixed. The resulting concentration-response curves (**Figure 1b**) unambiguously show a shift to the left in the response, i.e. an equal cAMP production can be achieved with a stimulus of less than half the 1 g control, as the EC_50_ values shift from 17.7 nM to 4.65 nM in hypergravity. As we did not observe any change in Emax, data were normalised between 0 (no response) and 1 (maximal response).

**Figure 1:**
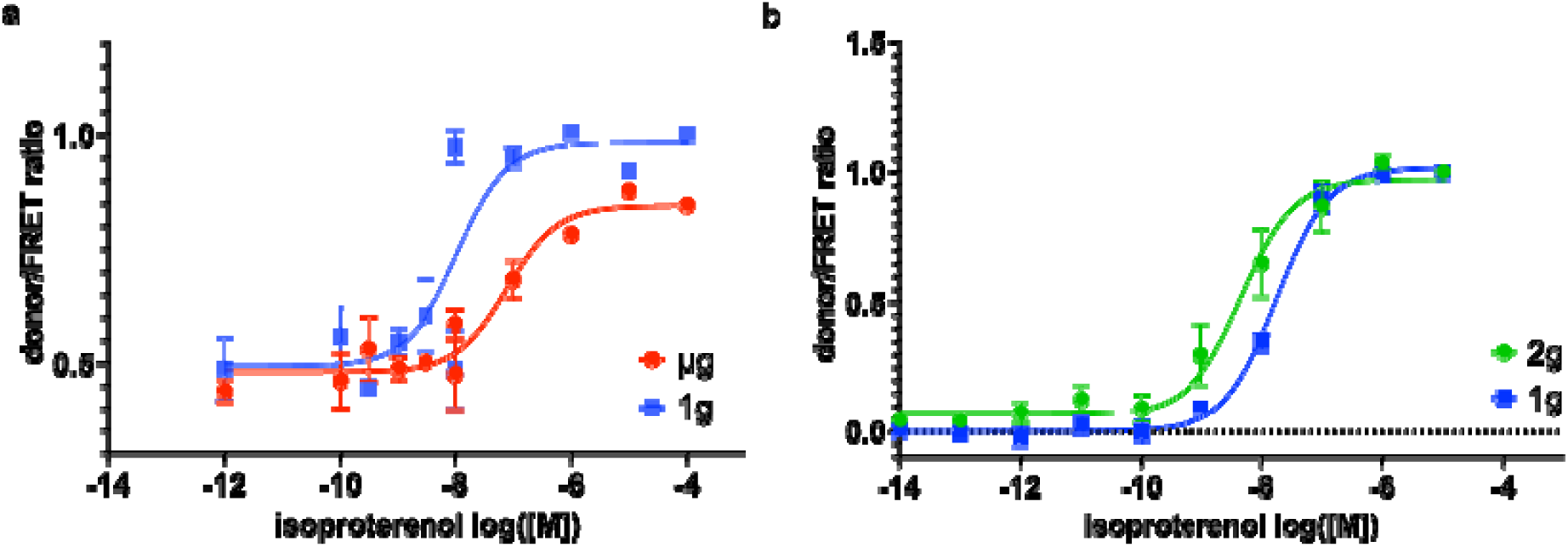
Effect of altered gravity on the isoprenaline-induced production of the second messenger cAMP, mediated by β-AR receptors. (a) Concentration response curve in HEK293 stimulated with increasing concentrations of isoproterenol while exposed to microgravity in a clinostat (µg) or ground control conditions (1g). cAMP levels are measured using the EPAC-S^H187^ FRET biosensor; the ratio between donor to acceptor fluorescence of each cell was measured in an epifluorescence microscope and plotted upon normalization to saturating isoproterenol stimulus. Each concentration point was obtained by averaging FRET ratios from 2 experimental sessions with at least 60 individual cells per concentration. Each series was repeated twice. Data are fit to an agonist vs response curve with Hill slope fixed to 1, yielding EC_50_^control^=10.6 nM (logEC50 95%CI[−8.642 to −7.242]) and EC_50_^µg^=78.5 nM (logEC50 95%CI[−7.657 to - 6.546]). (b) Concentration response curve in HEK293 stimulated with isoproterenol while exposed to hypergravity (2g) in a centrifuge or control conditions (1g). Cells, transfected as in (a), were grown in 96 well plates, and fluorescence readouts measured in microplate reader. Concentration response curves were normalised and data from n=6 experimental sessions was averaged. Fitting yields EC_50_^control^=17.7 nM (logEC50 95%CI[−7.861 to −7.644]) and EC_50_^2g^=4.65 nM (logEC50 95%CI[−8.7 to - 7.996])). Mean and SEM are indicated.

We would like to note here that albeit the two set of experiments depicted in Figure 1 were conducted in different experimental geometries, namely FRET ratios from single cells (µg) vs plate averages (hypergravity) further measured on two different set of equipment (microscope for µg vs microplate reader for 2g experiment), the 1g control readouts are in excellent agreement.

In order to understand better the molecular determinants of the steady state behavior, we moved to conduct kinetic measurements of cAMP production in response to an adrenergic stimulus in HEK293 cells, comparing cells exposed to altered gravity to ground control samples. To achieve this objective, it was necessary to exploit an experimental configuration allowing real-time microscopy imaging while the cells were exposed to altered gravity conditions. Therefore, we used the Hyperscope, a centrifuge-mounted microscope available at the Deutsche Luft und Raum Zentrum in Cologne (See **Material and Methods**). Using the Hyperscope we observed real-time cAMP accumulation in cells, transfected with the FRET biosensor and exposed to 2g hypergravity conditions, as they were perfused with the β-AR agonist isoproterenol. As soon as the agonist is added, using a peristaltic pump allowing exchanging medium in the imaging chamber with medium containing 1 µM isoproterenol, an increase in donor emission mirrored by a drop in FRET signal can be observed (**Figure 2a**). More snapshots of this activation are displayed in **Figure S2.**

**Figure 2:**
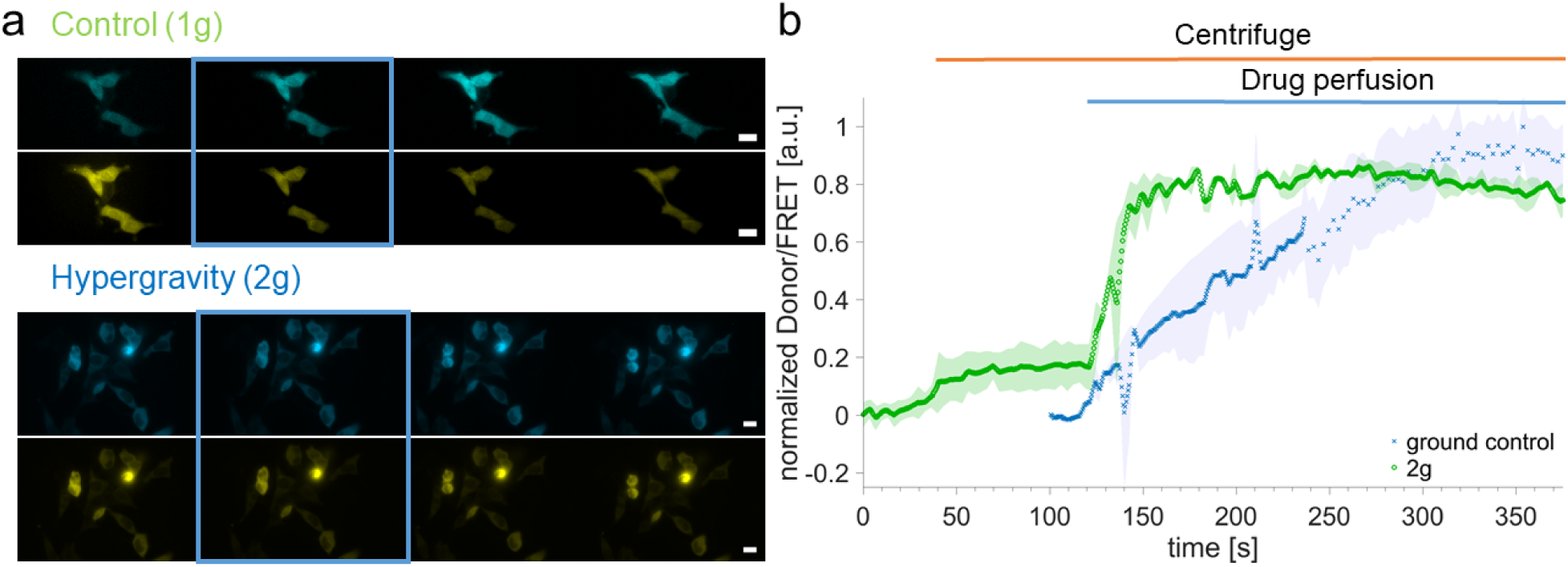
Kinetic of β-adrenergic cAMP production in single cells under control and hypergravity conditions in HEK293 cells. HEK293 cells transiently transfected with EPAC-S^H187^ were imaged in the Hyperscope, an epifluorescence microscope mounted on a centrifuge able to generate 2g conditions, or in the reference frame of the laboratory. (a) Representative snapshots from the donor and FRET microscopy channels at different time points before stimulus (1 µM isoproterenol), shortly after (rising phase), at maximal response, and after maximal response. Blue frames indicate the addition of the drug. Scale bars are 20 µm. (b) The normalized donor to FRET ratio was plotted as a function of time. Traces are the average of n=2 experiments for both hypergravity (green, 34 cells) and control conditions (blue, 12 cells). For the hypergravity trace (green circles) the centrifuge was activated at t=40s, and perfusion with 1 µM isoproterenol was initiated at t=120s. Shading represent SEM.

**Figure 2b** illustrates that when cells are exposed to 2g conditions in the Hyperscope, cAMP accumulation as reported by the donor/FRET ratio risemore rapidly than in the control conditions. In these experiments, the centrifuge was activated at approximately 40 s from the beginning of the experiment (orange line), and then perfusion of 1 µM isoproterenol was started at 120 s from the start of the experiment using a peristaltic pump (see **Materials and Methods**). The control experiment has a shorter baseline since no centrifuge activation time is required, **although** the measurements were conducted on the same microscope in the same experimental setting as the 2g measures. The data further show a small constitutive increase in cAMP as soon as the centrifuge is switched on and in the absence of any ligand, a form of constitutive activity which may relate to a mechanosensitive response of the cells to the increased gravity (Cullum et al. 2024; Sirbu et al. 2024).

Having unambiguously shown that β-AR downstream signalling is affected by altered gravity, next, we addressed the question of whether hypergravity conditions directly or indirectly, e.g. by a rearrangement of plasma membrane biophysical context, affected the β-ARs responsiveness. G protein-coupled receptors can switch their conformation to accommodate a ligand on the extracellular side, and/or a G protein on the intracellular side (De Lean, Stadel, and Lefkowitz 1980). The concomitant interaction with both a ligand and a G protein can lock the receptors in an active state, which supports nucleotide exchange at the G protein and thereby its activation. There may be instances which favor this occurrence, i.e. where the receptor conformation is altered by its surroundings.

In order to determine if in 2g the GPCRs were more prone to reside in an active conformation, we employed a conformational biosensor based on a circularly permutated green fluorescent protein (cpGFP) inserted in the third intracellular loop of the β2-AR (Patriarchi et al. 2018), within the construct β2-AR-cpGFP. In this instance, however, we did not observe any change in conformation of the receptor in response to the altered gravity stimulation (**Figure 3 & Figure S3**). This suggests that the molecular changes yielding the difference in the kinetic response displayed in **Figure 2** are to be found downstream of receptor activation.

**Figure 3:**
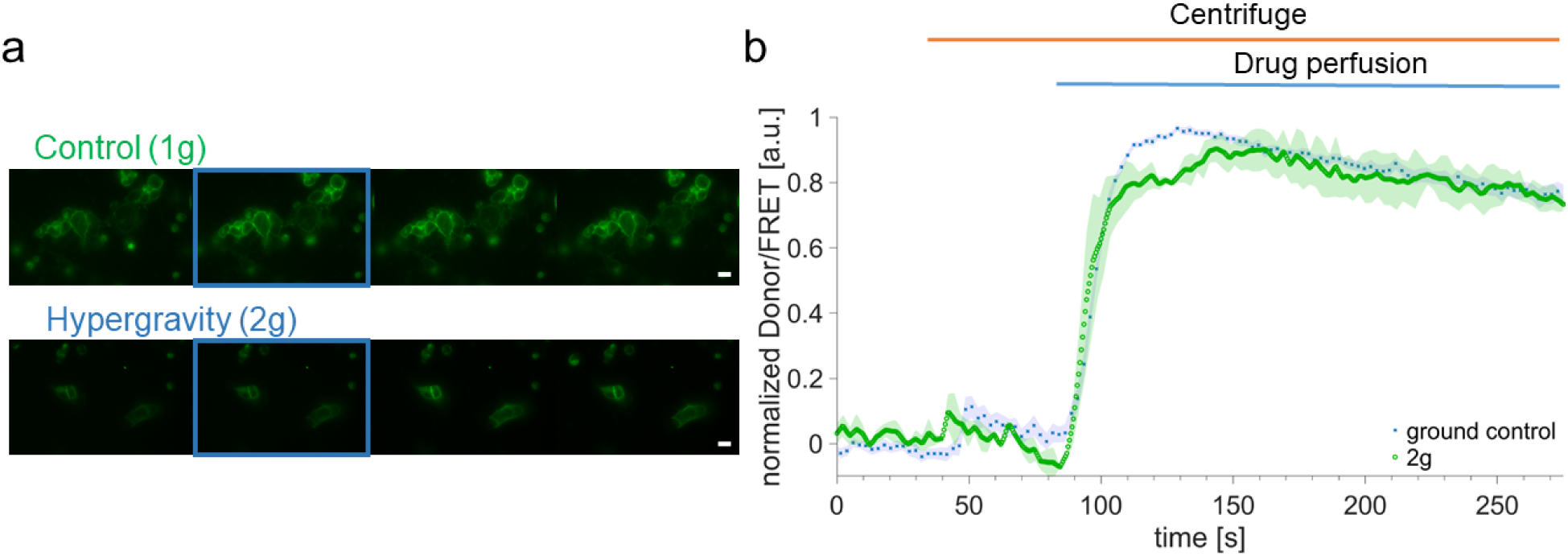
β2-adrenergic receptor activation kinetic in HEK293 cells under control and hypergravity conditions. (a) Representative image series at different time points before stimulus, shortly after (rising phase), at peak, and after peak. Blue frames indicate the immediate biosensor response. Scale bars are 20 µm. (b) Fluorescent intensity of the β2-AR-cpGFP biosensor reports receptor activation with 1 µM Isoproterenol. Averaged traces represent ground control (blue, n=1 experiments, 20 cells) and 2g (green, n=2 experiments, 14 cells). Shading represents SEM.

Before turning to a more detailed investigation of the possible molecular mechanisms giving rise to effects observed in Figures 1 and 2, we repeated our measurement in a cell line where β-ARs play an important physiological role. In heart muscle cells, cardiomyocytes, β-ARs contribute to regulate contractility, and therefore we investigated the effects of hypergravity on the activation kinetics of cAMP in H9c2 cells, a rat-derived cardiomyocyte-like cell line (Kuznetsov et al. 2015). In **Figure 4a** we see representative image series of the activation of H9c2 by measuring the activation with the FRET biosensor EPAC-S^H187^. We observe an increased signal in the cyan donor channel whereas the signal decreases in the yellow FRET channel after activation. More snapshots are shown in **Figure S4. Figure 4b** illustrates the traces of the extracted, corrected and normalized ratio of the two signals for 1g control cells and cells exposed to hypergravity. In line with the observation in HEK293 cells, control H9c2 cells (1g) display an apparently slower rate of cAMP accumulation than hypergravity cells upon perfusion with isoproterenol. Upon treatment of H9c2 cells with retinoic acid, a driver of cardiac maturation for this cell line, cAMP production kinetics (**Figure S5**) appear to happen on a faster scale than shown in **Figure 4**, albeit the limited statistics of the experiments conducted under these conditions do not allow to draw firm conclusions. Although these results point to potential cell-type specific differences in the kinetic of cAMP accumulation, they consistently show an increased cAMP production rate in hypergravity conditions.

**Figure 4:**
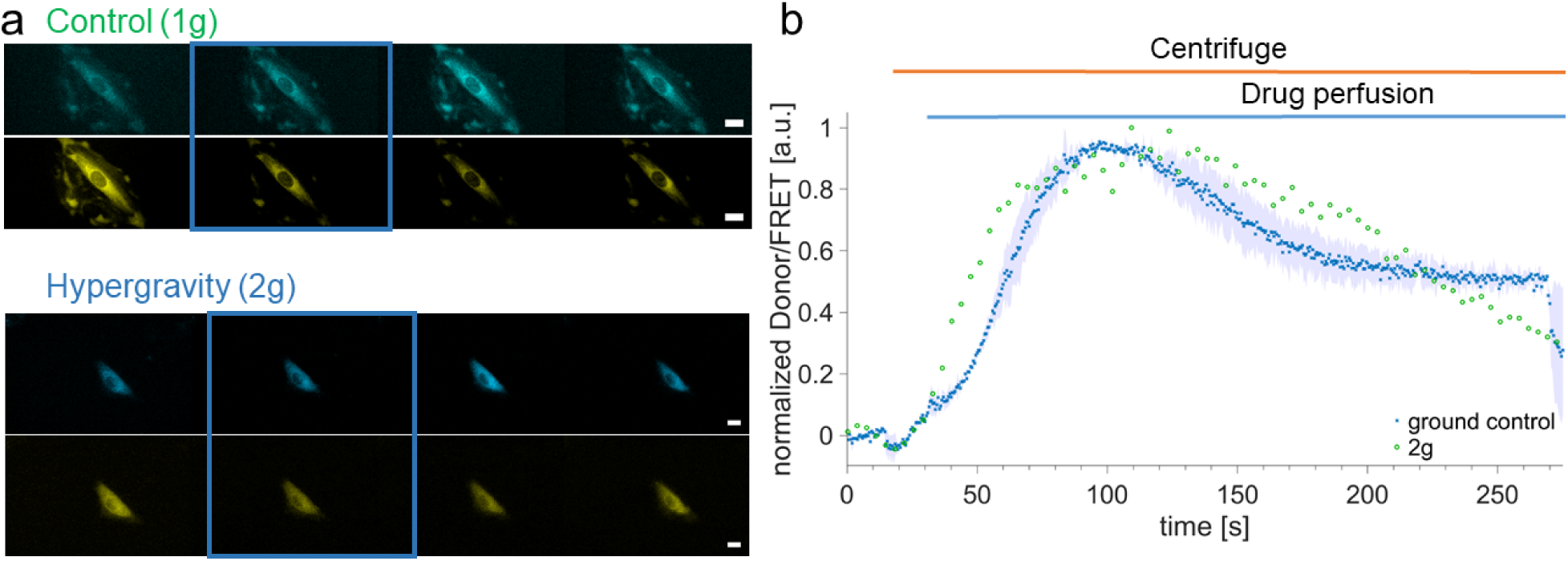
Kinetic of β-adrenergic cAMP production in single cells under control and hypergravity conditions in H9c2 cells. (a) Representative image series at different time points before stimulus, shortly after (rising phase), at peak, and after peak. Frames indicate the first change in channels fluorescence. Scale bar is 20 µm. (b) Averaged traces represent control (blue, n=2 experiments) and 2g (green, n=1 experiment) via FRET readout with the EPAC-S^H187^ sensor after 1 µM stimulation with Isoproterenol. Error bands represent SEM

Based on 1g data, we and others have observed an impact of cytoskeletal organization and receptor coupling on subsequent signaling and trafficking of G protein-coupled receptors (Scarselli, Annibale, and Radenovic 2012). For instance, in our hands an enhanced polymerization of the F-actin, using the actin mutant S14C (**Figure 5a**), leads to a blunting of receptor signaling, as seen by a shift to the right of the EC50 of cAMP concentration-response curves (**Figure 5b,c**).

**Figure 5:**
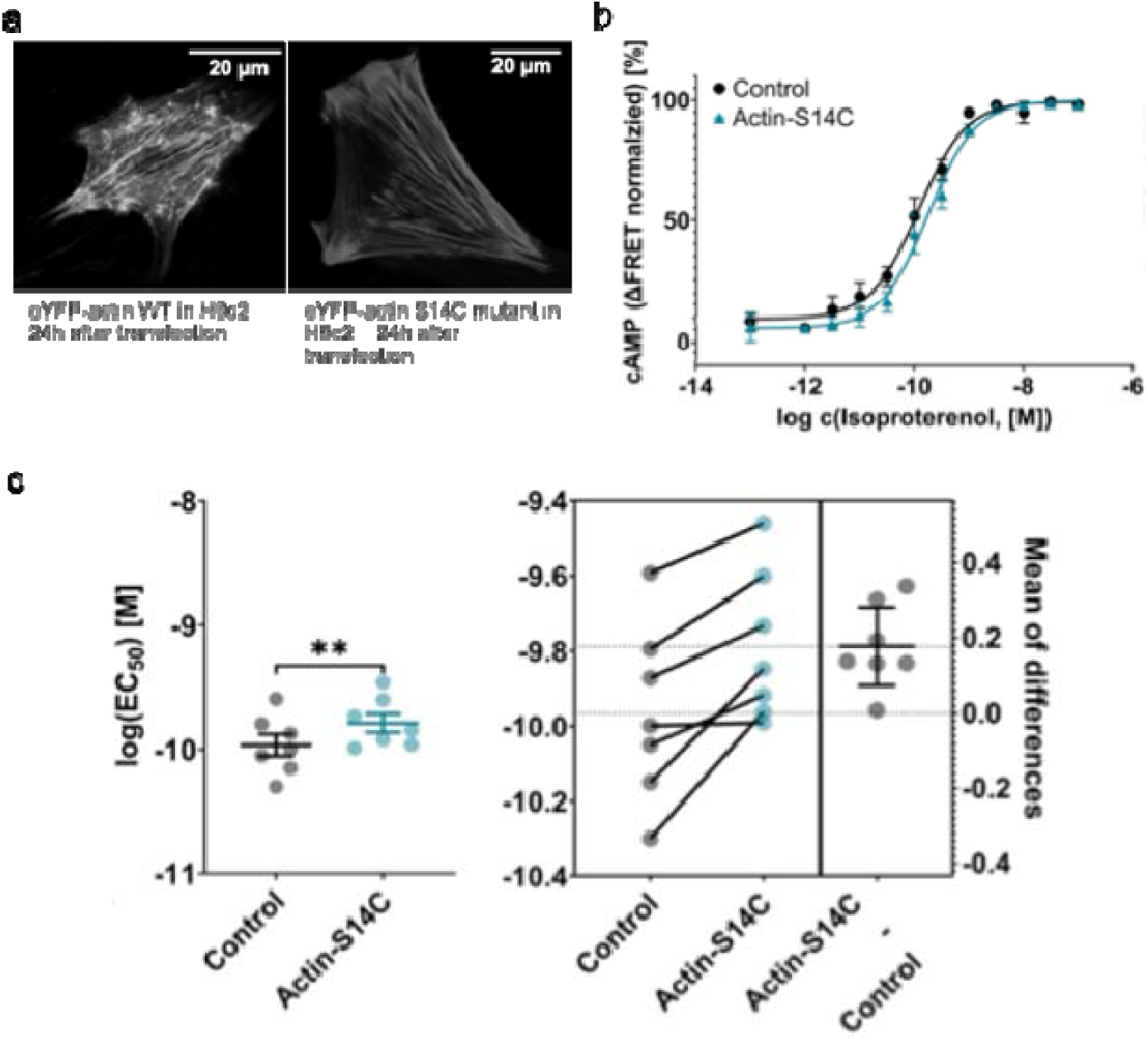
Effects of cytoskeletal remodeling (actin hyperpolarization) on β2-AR downstream signaling. a) confocal images of H9c2 cells transfected with eYFP-actin or eYFP actin S14C mutants, displaying the hallmark hyperpolarized cytoskeleton and increase in stress fibers . b) Average cAMP concentration response curve measured from HEK293 cells expressing the EPAC-cAMPH^187S.^ c) Individual EC50 values from n=7 independent experiments paired in order to display an increase in EC50 (loss of response) in cells where actin is hyperpolymerised.

This led us to hypothesize that the observed changes in the cAMP downstream signaling caused by altered gravity conditions could be underpinned by morphological changes in the actin structure of the cytoskeleton.

To investigate the influence of altered gravity conditions onto the actin skeleton we labeled HEK293 cells with lifeact-EGFP and exposed cells to simulated microgravity for 24h, matching the experimental conditions employed in **Figure 1a**. After exposure cells were rapidly fixed and the nuclei were stained with Hoechst 33342 before confocal images were acquired. Representative images under normal gravity (1g) and simulated microgravity (µg) are displayed in Error! Reference source not found.**Figure 6a,b**. More images, organized in mosaics, are shown in **Figure S6** for HEK293 cells, do not indicate any qualitative difference between the two conditions, either at the basolateral membrane where cortical actin is present, or in the perinuclear cell region. In order to conduct a more quantitative assessment, we conducted an orientation analysis of cellular actin filaments. The result of this analysis, which starts with an identification and thresholding of the filaments, yields a histogram indicating the average orientation of the actin filaments in the cells (**Figure 6c-f**). **Figure 6c-d** shows the orientations collected from actin filaments imaged in over 40 cells at the basolateral membrane in 1g (**Figure 6c**) and µg conditions (**Figure 6d**). We could not identify a clear pattern of change between 1g and µg, suggesting that no quantifiable disorganization takes places upon 24h microgravity, at least at the level of the microfilaments detectable using a diffraction-limited imaging approach such as confocal microscopy. This would seem to argue against cortical actin being responsible for the signaling changes reported for adrenergic signaling in altered gravity settings. The same considerations apply for observations in a confocal plane through the middle of the cells (**Figure 6e-f)**.

**Figure 6:**
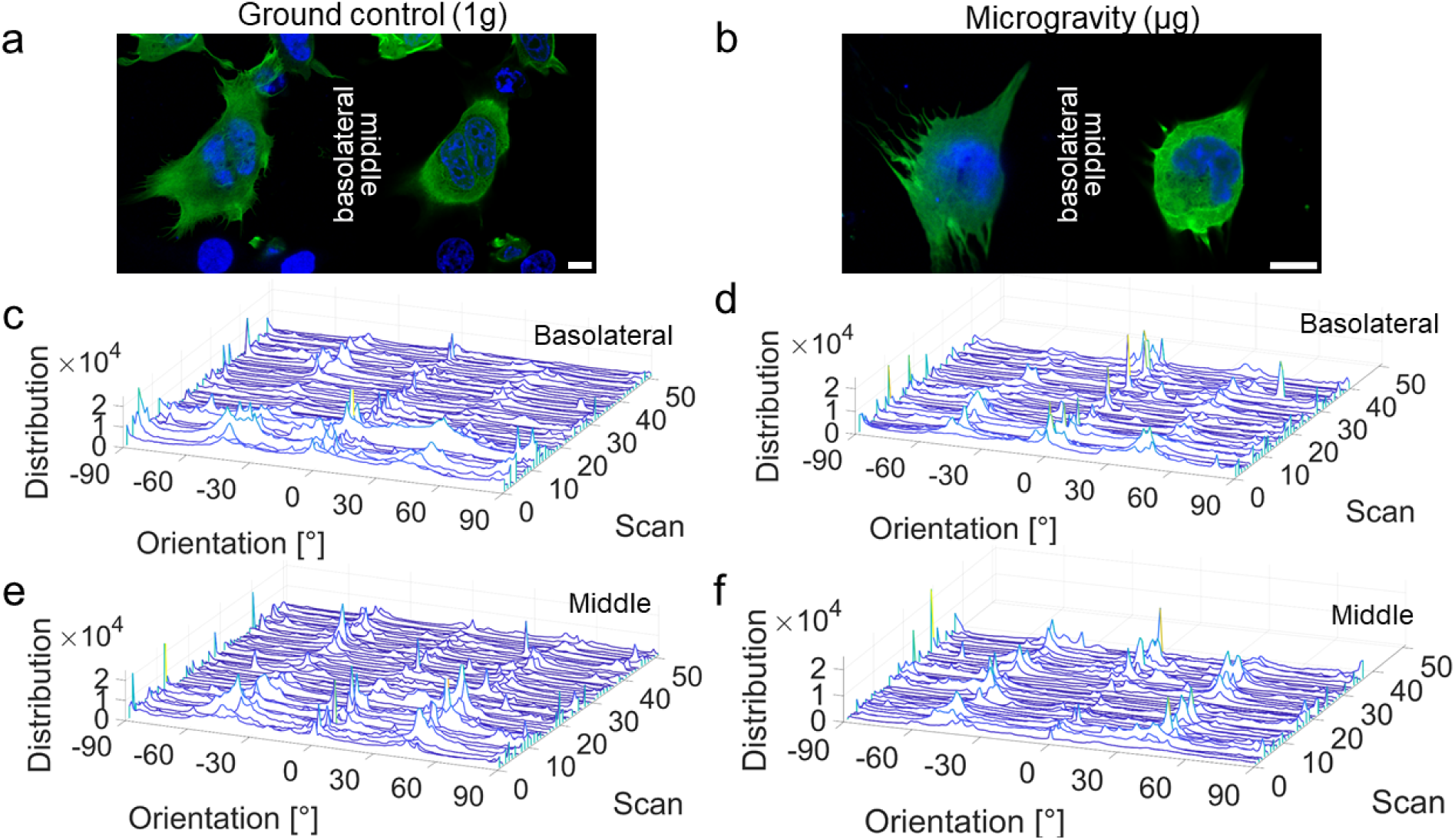
Effect of microgravity on the cytoskeletal actin of HEK293 cells. HEK 293 were transiently transfected with the fluorescently tagged actin label lifeact-EGFP and imaged after being exposed to microgravity conditions or the laboratory reference and subsequently fixed. Extraction of orientation analysis using OrientationJ for control and µg conditions at the basolateral membrane and a plane through the middle of the cell. a, b) Structural features were extracted by confocal microscopy, conducted both at the basolateral membrane and in the middle of the cell. Actin is shown in green (Lifeact-EGFP) and the nucleus was stained with Hoechst 33342 (blue). Scale bars are 10 µm. Histogram indicating the average orientation of the actin filaments at the basolateral membrane c) 1g, d) µg, and a perinuclear plane e) 1g, f) µg. Basolateral 0g: 40 cells, middle: 37 cells, 1g: basolateral and middle: 51 cells from 2 experimental sessions.

We could only observe changes in basic parameters like area, perimeter and roundness of cells between 1g and µg conditions (**Figure S7**), which do not specifically reflect changes to cytoskeletal arrangement, albeit indirectly confirming a cell-wide effect of the microgravity exposure. To rule out that overexpression of lifeact-EGFP, known to stabilize actin cytoskeletal integrity may have masked physiological changes, we also employed hiPS cells where an EGFP fusion was inserted heterozygously at the N-terminus of β-actin via Crispr/Cas9. Qualitative inspection did not display any significant changes in actin morphology between the two conditions also for this cell-type (**Figure S8**).

## Discussion

We measured the effects of altered gravity on the cAMP response to adrenergic stimulation in HEK293 cells, a well-established human cell line used broadly in in-vitro pharmacology studies. We found that simulated µg and hypergravity have opposing effects on the potency of adrenergic-mediated cAMP production. The observed increase in EC_50_ (loss of potency of the adrenergic stimulation) in simulated microgravity (**Figure 1a**) is in line with the reduced cAMP dependent PKA activity observed for rats from SpaceLab-3 reported by Medniekis et al. (Mednieks et al. 1987). At the same time, Convertino et al (Convertino et al. 1997) and Barbe et al (Barbe et al. 1999) (Albi et al. 2011) recently studied the cAMP response to TSH Receptor stimulation in a sounding rocket experiment (6 minutes of microgravity) and observed a decrease in cAMP levels, hypothesizing a reorganization of membrane microdomains as the mechanistic basis for this decrease. The observed decrease of EC_50_ in hypergravity (**Figure 1b**) suggests increased responsiveness of cells to increased gravity conditions (increased potency of adrenergic stimulation), which would align with recent results obtained by Acharya et al (Acharya et al. 2019): here, hiPS-derived cardiomyocytes were subjected to an increasing number of parabolas, which comprise intervals of hyper- and simulated microgravity, and found increased responsiveness to isoproterenol as the number of parabolas increased. However, given the combination of two opposing gravity conditions in each parabola, it is hard to establish a direct parallel between these and our data.

Our steady state observations are mirrored by an enhanced kinetics of cAMP accumulation in the HEK293 cells subjected to hypergravity conditions, which is also observed in a more physiological cardiac model, namely H9c2 cells (Hescheler et al. 1991) (**Figure 4**). These observations, conducted by live cell imaging of cells exposed to 2g conditions, also indicate a mild increase in receptor constitutive activity (**Figure 2b**) once 2g conditions are achieved, before any agonist is added. These results may be reconciled with a degree of mechanosensation of the adrenergic receptors, as recently highlighted by several groups (Cullum et al. 2024; Sirbu et al. 2024; Virion et al. 2019), which would be triggered by the increased force felt in 2g conditions. However, quite interestingly, our measurements did not show any effect in terms of conformational changes at the β2-AR, the predominant adrenergic receptor expressed in HEK cells (**Figure 3**). It is possible that conformational rearrangements affect domains which do not alter tension on the cpGFP sensor used in our constructs, or that altogether other changes in the downstream signaling pathway may instead be involved.

The actin cytoskeleton, in particular the portion that underlies the cell membrane, has crucial functions in regulating the plethora of membrane receptors that populate the plasma membrane and act as the gateway to the intracellular biochemistry (Stelzer 2021; Boltz et al. 2022; Kockelkoren et al. 2024; Scarselli, Annibale, and Radenovic 2012). For this reason, we investigated whether altered gravity may affect the arrangement of cortical actin and thereby affect the organization and function of the membrane receptors. Our data (**Figure 5)** indicate that increased actin polymerization reflects decreased potency for adrenergic signaling in HEK293 cells.

Since the very onset of real and simulated microgravity experiments, the cellular cytoskeleton has been one of the key organelles investigated using fluorescence and ultrastructural methods to monitor the effect of altered gravity (Ross 1984; Mednieks et al. 1987; Philpott 1990). Analogously to the human skeletal system, which suffers under extended microgravity exposure, it was expected that the polymer structures responsible for maintaining cell shape would be amongst the most affected by the loss of gravity. Within this context, actin integrity and organization in isolated cells subjected to altered gravity have been an important focus of research since early space research experiments. These experiments have ranged from plant cells to human cells, the latter including a variety of model systems, encompassing immune cells, cancer cells and cardiac cells, to name only a few (Nassef et al. 2019; Lin et al. 2016; Janmaleki et al. 2016; Nabavi et al. 2011; Yang et al. 2008; Versari et al. 2007; Rosner et al. 2006; Crawford-Young 2006; Blancaflor 2013; Siamwala et al. 2010).

We therefore tested whether sizable actin remodelling could be observed in our samples when exposed to simulated microgravity. However, in our model and under the altered gravity conditions we employed, we could not detect any quantifiable change to actin morphology, suggesting that other molecular mechanisms may be at play (**Figure 6**). We note here that despite the fact that our overall knowledge of cytoskeletal actin organization in cells in normal gravity is very developed (Svitkina 2018), there appears to be no univocal answer to the question as to exactly how the actin cytoskeleton is rearranged in altered gravity in general, and microgravity in particular, with reports pointing to disorganization, redistribution, reduction or no change at all with respect to the F-actin content in the cell (reviewed by (Crawford-Young 2006)).

In light of the importance of cAMP signaling in regulating cardiac function and many other physiological processes (Gancedo 2013) coupled with the possible increase in space exploration in the near future, it is important to understand how changes in gravity affect cAMP metabolism.

Our data indicate that altered gravity affects the cAMP cascade, yet the precise molecular switch remains elusive. We excluded two plausible mechanisms—direct conformational changes at the third loop of the receptor and actin cytoskeleton remodeling—but several other steps in the signaling pathway may contribute. Previous studies have implicated altered activity of adenylyl cyclases (ACs) and phosphodiesterases (PDEs), the enzymes synthesizing and degrading cAMP, respectively, and recent work supports altered PDE and AC gene expression under altered gravity (Mednieks et al. 1987; Sirbu et al. 2024; Mitrokhin et al. 2024). Our 24h endpoint experiments are consistent with this possibility, although changes were not uniform across PDE isoforms. Furthermore, comparative analysis of different GPCRs revealed that β2-adrenergic receptors and melanocortin-4 receptors (Figure S9) display distinct mechanosensitive behaviors, pointing to receptor-specific responses and suggesting that the relevant switch resides at the plasma membrane. Although our cpGFP-based conformational biosensor did not reveal sizable changes at the β2-AR (Figure 3), this negative result cannot exclude a mechanosensitive contribution of this receptor. As recently reviewed (Shetty, Sirbu, and Annibale, 2025), the β2-AR has been implicated in mechanosensitive processes, and movements of Helix 8—thought to underlie such effects—may not be detected with the biosensor employed here. This interpretation is consistent with our recent findings on osmotic swelling at the β2-AR (Sirbu et al., 2024), further suggesting that the choice of readout strongly influences the detection of receptor mechanosensitivity. Altogether, this highlights the complexity of gravity-induced modulation of cAMP signaling, involving multiple checkpoints rather than a single switch. Future studies dissecting these pathways in detail will be essential to predict and manage pharmacological responses during spaceflight.

## Methods

### Altered gravity experiments

All altered gravity experiments were performed at the German Aerospace Centre (DLR, Köln, Germany) using the following equipment:

| Product Name | Manufacturer |  |
| --- | --- | --- |
| Multi Sample Incubator Centrifuge (MuSIC) | German Aerospace Center, Institute of Aerospace Medicine, Department of Applied Aerospace Biology<br>Based on a standard cell incubator manufactured by Binder, Germany | • Hypergravity Load on Sample: 1g - 50g<br>• Acceleration ramp: up/down adjustable<br>• Exposure time: seconds – weeks<br>• Temperature range: RT up to 50 °C<br>• Humidity range: up to 95 % RH<br>• Equipped CO<sub>2</sub>-gas mixing nozzle<br>• Auto-sterilization via hot air at 180 °C<br>• Anti-condensation humidifying-system |
| Hyperscope Live-Cell Imaging Platform | Centrifuge manufacturer: AMST, Austria<br>Microscope manufacturer: Carl Zeiss, Jena, Germany<br>Hyperscope Platform and swing-out gondola: German Aerospace Center, Institute of Aerospace Medicine | • Zeiss Axio Observer Z1 live-cell imaging microscope<br>• Swing-Out Platform to Avoid Shear Forces on Samples<br>• DLR short-arm Human Centrifuge in :envihab facility<br>• Centrifuge Radius: 3,80 m<br>• Maximum radial acceleration: 4g<br>• Acceleration 0g – 4g in 30 s<br>• Experiment runtime: up to 24h<br>• Fully automated Microscope Setup<br>• Remote Control for every action also live possible during Centrifuge Operations<br>• Objectives: 10x air (NA 0.3), 20x air (NA 0.4), 25x oil (NA 0.8), 40x oil (NA 1.4), 100x oil (NA 1.4)<br>• LEDs: 385 nm, 475 nm, 555 nm, 590 nm, 630 nm<br>• Filters: DAPI, EGFP, Alexa 555 (Cy3), mCherry, Alexa 660 (Cy5), DAPI/EGFP/mCherry triple bandpass<br>• Camera: CMOS, 128fps, highly sensitive, time-lapse<br>• Online focus stabilization via Definite Focus.2<br>• PeCon incubation chamber for constant temperature and CO<sub>2</sub><br>• Administration of drugs or other substances while the experiment is running |
| Slide Flask Clinostat | German Aerospace Center, | • Simulated Microgravity of |
|  | Institute of Aerospace Medicine,<br>Department of Applied Aerospace<br>Biology | 0.008g<br>• Rotation speed: 60 rpm<br>• Exposure time: seconds – weeks<br>• Incubator Temperature range: RT to 42°C<br>• Incubator Humidity range: up to 95 % RH<br>• Incubator Equipped with CO<sub>2</sub>-gas mixing nozzle<br>• Incubator Auto-sterilization via hot air at 180 °C |

### Cell Culture

HEK293T cells (ECACC 96121229, Sigma-Aldrich) (henceforth HEK293) and H9c2 cells (ATCC - CRL-1446) were used throughout this work for hypergravity experiments using Multi-Sample Incubation Centrifuge (MuSIC) and the Hyperscope, as well as microgravity experiments using a clinostat. Cells were grown in Dulbecco’s Modified Eagle’s Medium (DMEM, Pan Biotech), supplemented with 2 mM L-glutamine (Pan Biotech), 10 % (v:v) heat-inactivated Fetal Calf Serum (Biochrome), 100 μg/mL streptomycin, and 100 U/mL penicillin (Gibco) at 37°C in a 5 % CO_2_ incubator. HEK293 cells were grown in T75 cm^2^ flasks. Upon reaching approximately 80 % confluency, cells were washed with 5 mL phosphate-buffered saline without Ca^2+^ and Mg^2+^ ions (DPBS, Sigma-Aldrich), trypsinised using 3 mL 0.05 % / 0.02 % trypsin/ethylenediaminetetraacetic acid solution (Pan Biotech) and passaged every 2 to 3 days. Cells were routinely tested for mycoplasma infection using MycoAlert Mycoplasma Detection Kit (Lonza). hiPSCs (Coriell Institute, Allen Cell Collection, AICS 0016) were grown at 37 °C with 5% CO2 and 5% O_2_, whereas differentiated cultures were maintained at 5% CO_2_ and atmospheric (21%) O_2_. H9c2 maturation towards a more cardiomyocyte-like phenotype via retinoic acid was achieved according to a protocol described by Pereira, et al. (2011), in which serum was reduced to 1% and cells were supplemented daily with 10nM of RA (Sigma) for 11 days. For comparison, some cells were maintained in either 10% FBS (control), or in 1% FBS for the same duration, but without RA (Pereira et al. 2011).

### Cell preparation for plate reader acquisition following hypergravity experiments

HEK293 cells were seeded in T75 cell culture flasks and transfected prior to transport to the ground based facility (GBF). Transfection was conducted using JetPrime (Polyplus) according to the manufacturer’s instructions using the Epac-S^H187^. Epac-S^H187^ was a gift from Kees Jalink (Addgene plasmid # 170348 ; http://n2t.net/addgene:170348 ; RRID:Addgene_170348). Media for HEK293 cells was changed 4h after transfection. 24h post-transfection, cells were re-seeded into PDL-coated black-wall (TPP), black bottom 96-well plates at a density of 40,000 cells per well. Ground control plates were placed at 37 °C in a 5 % CO_2_ incubator while the hypergravity 2g plate samples were placed in the MuSIC, which was pre-set to generate a gravity of 2g @80 rpm, while maintaining cells at 37°C and 5% CO_2_ conditions. After hypergravity exposure cells were immediately stimulated for 5 mins with 10 μL of 10-fold Isoproterenol (Isoproterenol hydrochloride, Sigma) solution applied to each well. Selected wells were stimulated with Forskolin (Sigma) and 3-isobutyl-1-methylxanthine (IBMX) (Sigma) to achieve maximal cAMP response for normalization purposes. Afterwards, cells were washed and medium was changed to HBSS buffer. Then, cells were fixed with a 4% Paraformaldehyde PBS solution supplemented with 0.2% Glutaraldehyde. Twenty minutes post-fixation, cells were washed with PBS and kept at 4°C until plate reader imaging. The ground control plates were also treated simultaneously with their respective 2g counterparts.

### Cell preparation for microscopic imaging following clinostat experiments

HEK293 and H9c2 cells were seeded in T75 cell culture flasks and transfected prior to transport to the ground based facility. There they were passaged into glass-bottom µ-slides (Ibidi) coated with Poly-L-Lysine (PLL), to be exposed to altered gravity the next day (48h post transfection). Transfection was conducted using Lipofectamine 2000 (Thermo Fisher Scientific) according to the manufacturer’s instructions using the Epac-S-H187, or mEGFP-Lifeact-7. mEGFP-Lifeact-7 was a gift from Michael Davidson (Addgene plasmid # 54610; http://n2t.net/addgene:54610; RRID:Addgene_54610). One to two days post transfection cells were exposed to the clinostat and treated with ligand (where applicable) before fixation. hiPSC-CMs were detached as single cells and seeded in Geltrex-coated, glass-bottom µ-slides (Ibidi, Gräfelfing, Germany). Generally, until the day of experiment hiPSC-CMs were cultured in a maintenance medium with 3 µM CHIR99021 and 1x RevitaCell supplement (Life Technologies, Carlsbad, CA, USA). A 2D rotation clinostat was used to simulate microgravity in vitro. µ-slides were attached in holders to the rotation bar. The clinostat had four bars which were simultaneously rotating allowing to load multiple µ-slides. The clinostat was placed inside a cell incubator to allow the cells to be kept at 37 °C and 5% CO_2_. 1g controls were placed on top of clinostat to be exposed to the same slight movements, apart from clinorotation, as cells exposed to µg. Cells were exposed to µg for 24h, quickly removed from the clinostat and stimulated where relevant with drugs by replacing the entire medium in the channel with media containing ligands at the appropriate concentration for a duration of 5 minutes. Then, cells were fixed with a 4% Paraformaldehyde PBS solution supplemented with 0.2% Glutaraldehyde. Twenty minutes post-fixation, cells were washed with PBS and kept at 4°C until microscopic imaging at a later time point.

### Hyperscope

The kinetic hypergravity experiments were performed on the *Hyperscope*, a widefield microscope that is mounted onto the DLR’s short-arm human centrifuge, which co-rotates together with the sample and experiences the same g-forces. The Hyperscope is a system that integrates the Axio Observer Z.1 live-cell fluorescence microscope mounted on a swing-out platform attached to one arm of the short-arm human centrifuge (SAHC1) at the DLR :envihab facility. This setup allows live-cell imaging of biological samples under hypergravity conditions of up to 2g. All settings of the microscope can be remotely controlled during centrifuge operation. The swing-out platform ensures that the gravity vector acts perpendicular to the samples, thereby avoiding unwanted shear forces (Lichterfeld 2024). Imaging settings for the FRET experiments were LED excitation at 385 nm, and collection of donor (mTurquoise2) emission between 475-523nm, and collection of FRET signal (mVenus) between 500-540 nm, by alternating filtercubes within the motorised turret of the Axio Observer Z.1. For the cpGFP biosensor experiments excitation was provied at 488 nm and collection between 500-550 nm. While the cells were exposed to 2g the microscope was controlled remotely from the outside. A perfusion system with a peristaltic pump (Masterflex® Ismatec® Reglo ICC Digital Pump), also remotely controlled, was connected using luer connectors to the µ-slides inlet and outlet. This setup allowed to add media with defined concentration of ligands at the desired time after the centrifuge startup. Cells were maintained at 37°C thanks to a stage insert incubator. Ground control experiments were ran with the exact same settings as those employed for the 2g experiments.

### Fluorescence plate reader analysis

Plate reader experiments were performed using a Synergy Neo2 plate reader (BioTek) equipped with a monochromator and filter optics. Expression levels of biosensor were measured with monochromator optics. For expression of biosensor the excitation was at 500/20 nm, and emission was at 539/20 nm. For FRET measurements, a range of CFP/YFP monochromators were used, which was excited at 430/20 nm, while 491/30 nm and 541/20 nm were used for emission.

### Confocal Microscopy

Images of actin structures in both HEK293 as well as hiPS cells were acquired on a confocal laser scanning microscope, Leica TCS SP8 and the same imaging conditions were applied throughout. All measurements were conducted with an HC PLAP CS2 40×1.3 numerical aperture (NA) oil immersion objective (Leica). The pixel size was 50 nm and the autofocus of the microscope was enabled during the acquisition to stabilize the focal position. EGFP was excited at 488 nm with a laser power of 1% and Hoechst 33342 was excited at 405 nm with a laser power of 2%. Emission of sequential scans was detected on hybrid detectors in photon-counting mode detecting in the range of either 420 to 479 nm (Hoechst 33342) or 492 to 548 nm (EGFP).

### FRET microscopy

HEK293 and H9c2 cells transiently transfected with Epac-S-H187 sensor were seeded in Ibidi µ-slides exposed to gravity conditions and stimulated followed by 4% Paraformaldehyde (PFA) with 0.2% Glutaraldehyde fixation. Cells were imaged at room temperature. An inverted microscope (DMIRE2, Leica Microsystems), equipped with an 40x HCX PL APO, 1.25 numerical aperture (NA) objective (Leica Micro-systems), dichroic beamsplitter T510lpxrxt (Thorlabs), Cool LED (pE-400 Series) and an Andor iXon Ultra EMCCD camera with a dual image splitter (OptoSplit II, Cairn Research), was used. We applied an excitation wavelength of 450 nm and collected fluorescence emission simultaneously at 470/24 nm and 535/30nm. Images were obtained at 500 ms exposure per frame. For live cell sequences movies were acquired for the number of frames needed for the cAMP concentrations to equilibrate. Fiji was used to extract fluorescence intensity values from single cells, which were corrected for back-ground and used to calculate FRET/ mTurquoise2 ratio.

### Data Analysis

For plate reader experiments, change in FRET for each well was initially calculated using Microsoft Excel 2019. For dose response curves from clinostat experiments and centrifuges data was fit in GraphPad Prism (v 9.5) to the following curve:

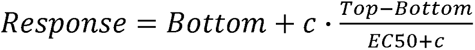

In which Top and Bottom are the plateaus in units of Response, c is the concentration of the agonist and EC50 is the concentration of agonist that gives a response halfway between Top and Bottom.

## Data availability

All relevant data are available from the corresponding author upon reasonable request.

## Code availability

All data analysis routines are available from the corresponding author upon reasonable request.

## Author Contributions

M.B.-P. Marc Bathe-Peters

I.S. Iqra Sohail

A.S. Alexei Sirbu

K.S. Katharina Schneider

T.P. Tommaso Patriarchi

A.A. Anantha Anilkumar

Y.L. Yannick Lichterfeld

C.L. Christian Liemersdorf

P.deL. Primal de Lanerolle

P.A. Paolo Annibale

I.S., M.B-P., P.A., A.S., K.S., A.A. acquired data. M.B.P., P.A., I.S., A.S., A.A analyzed data. T.P. and P.De.L provided materials and reagents. Y.L. and C.L. provided onsite technical support. P.A., M.B.P., P.deL. wrote the manuscript with input from all authors.

P.A. supervised the project. P.A. acquired funding.

## Acknowledgments

We are grateful to Heike Biebermann (Charite Universitätsmedizin, Berlin) for support during sample preparation. We are grateful to Leye Cooker (Cellbox gmbh) for support related to cell transport from St Andrews to Cologne.

## Ethics declaration

The authors declare no competing interests.

## Supplementary Figures

**Figure S1.**
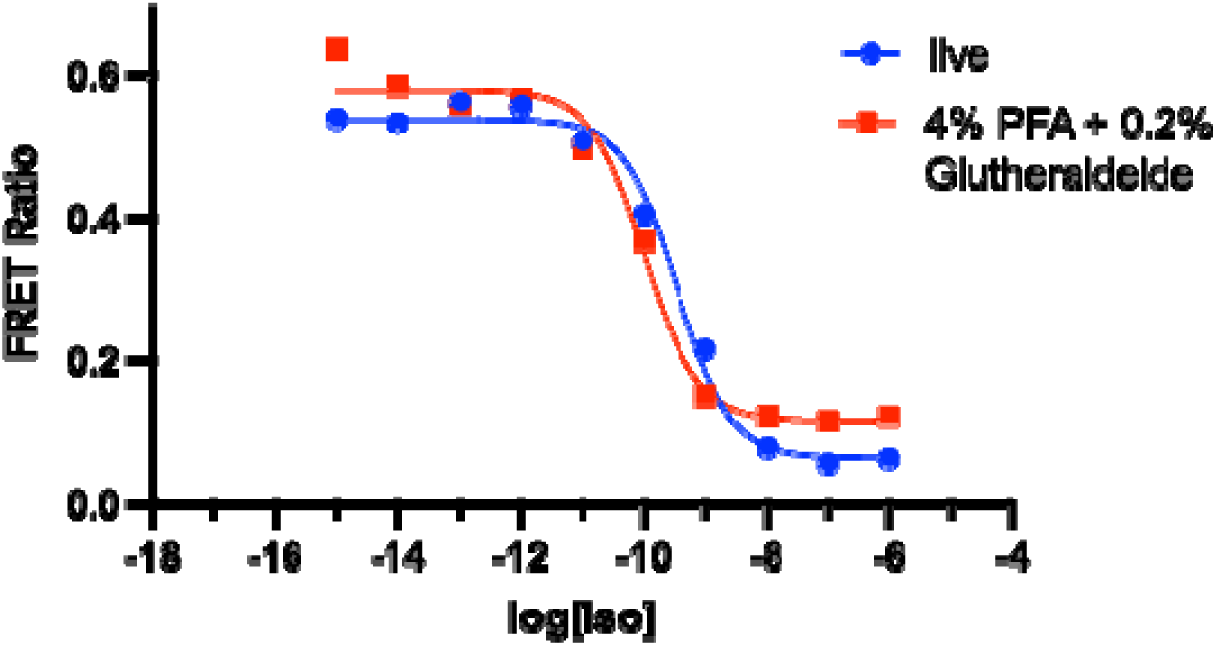
Effect of fixation on FRET readout for HEK293 cells transiently transfected with EPAC-S^H187^. cAMP concentration as a function of Isoproterenol(Iso) stimulus comparing live vs chemically fixed HEK293 cells using. Fixation was achieved by a combination of 4% Paraformaldehyde and 0.2% Gluteraldehyde, for 10 minutes at room temperature. n=1 replicate.

**Figure S2:**
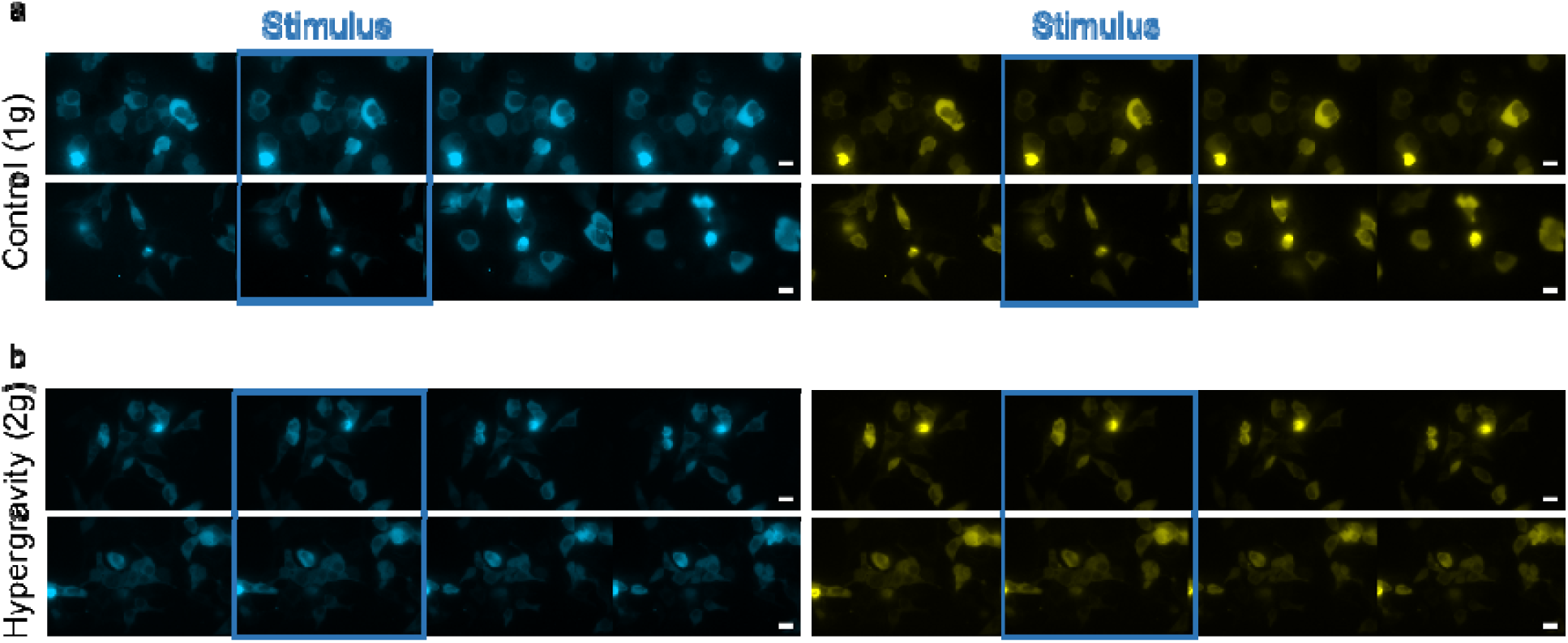
β-adrenergic activation measured by live FRET microscopy of HEK293 transfected the EPAC-SH187 sensor and exposed to 1g or 2g conditions in the Hyperscope. a) donor (mTurquoise2, left)and FRET (mVenus, right) signals from two representative experiments (out of n=5 experiments) where HEK293 cells were stimulated at the indicated time with 1 µM isoproterenol in 1 g conditions. b) donor (left)and FRET (right) signals from two representative experiments (from n=5 experiments) where HEK293 cells were stimulated at the indicated time with 1 µM isoproterenol in 2 g conditions. Each time sequence contains a snapshot stimulus, shortly after (rising phase), at peak response, and after peak. Scale bar is 20 µm throughout.

**Figure S3:**
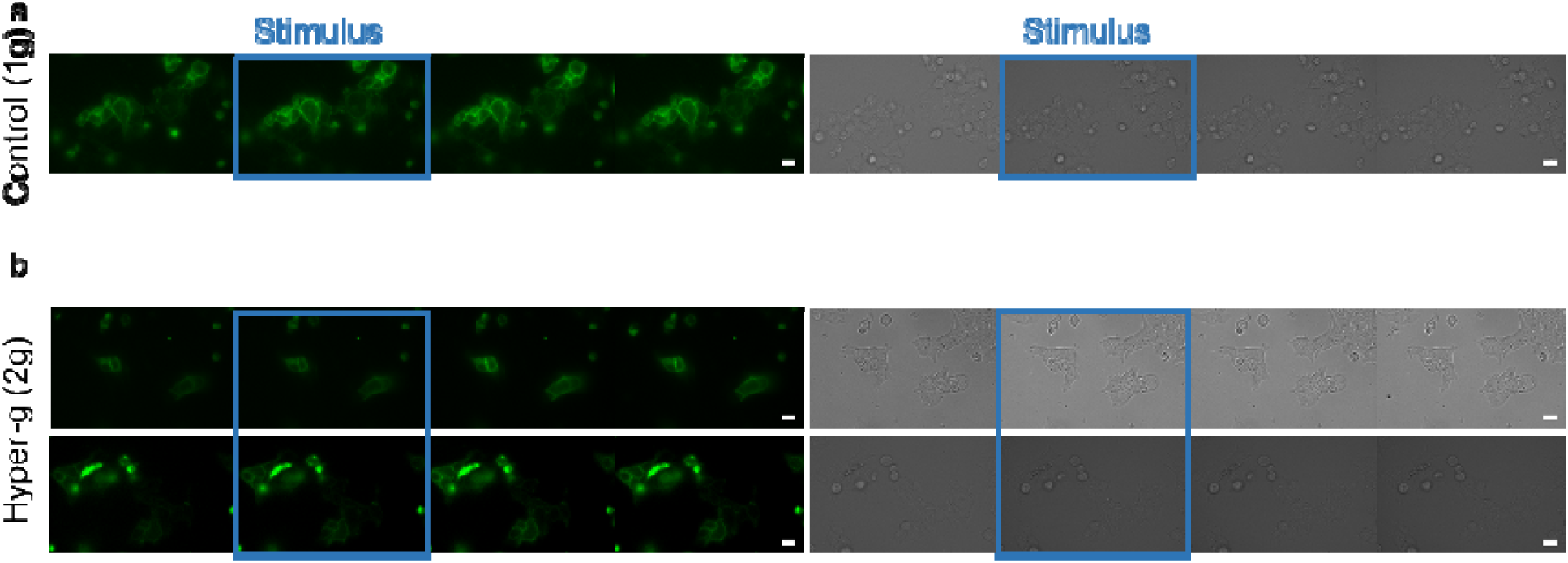
β-adrenergic activation measured by live FRET microscopy of HEK293 transfected the β2-AR-cpGFP sensor and exposed to 1g or 2g conditions in the Hyperscope. a) cpGFP (green, left) and DIC (gray, right) signals from a representative experiments where HEK293 cells were stimulated at the indicated time with 1 µM isoproterenol in 1 g conditions. n=1 experiment. b) signals from n=2 experiments where HEK293 cells were stimulated at the indicated time with 1 µM isoproterenol in 2 g conditions. Each time sequence contains a snapshot stimulus, shortly after (rising phase), at peak response, and after peak. Scale bar is 20 µm.

**Figure S4:**
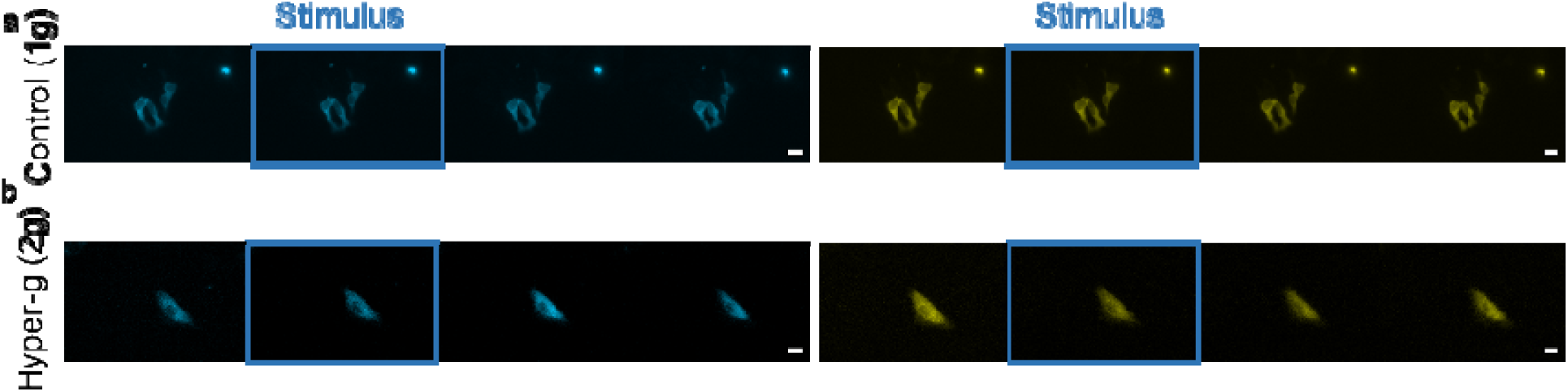
β-adrenergic activation measured by live FRET microscopy of H9c2 transfected the EPAC-SH187 sensor and exposed to 1g or 2g conditions in the Hyperscope. a) representative donor (mTurquoise2, left)and FRET (mVenus, right) signals from n=5 experiments where H9c2 cells were stimulated at the indicated time with 1 µM isoproterenol in 1 g conditions. a) representative donor (left)and FRET (right) signals from n=2 experiments where H9c2 cells were stimulated at the indicated time with 1 µM isoproterenol in 2 g conditions. Each time sequence contains a snapshot stimulus, shortly after (rising phase), at peak response, and after peak. Scale bar is 20 µm. Scale bar is 20 µm throughout.

**Figure S5:**
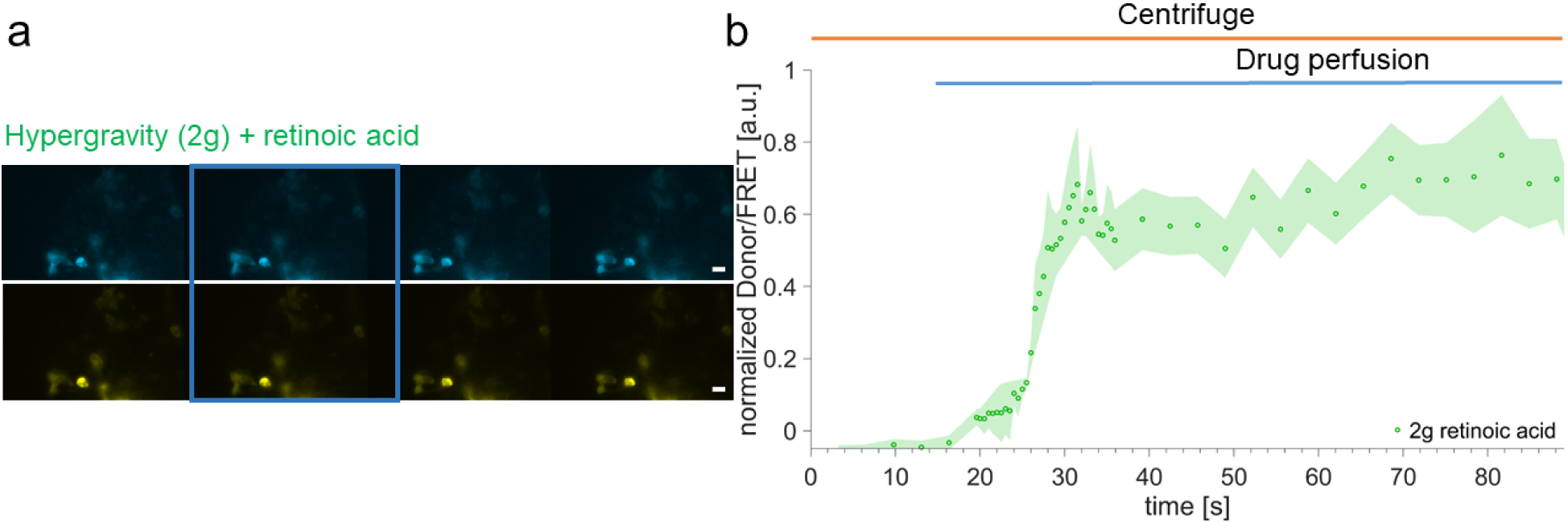
β-adrenergic activation in retinoic-acid treated H9c2 cells under hyper-g. (a) Representative image series at different time points before stimulus, shortly after (rising phase), at peak, and after peak. Frames indicate the first change in channels fluorescence. Scale bar is 20 µm. (b) Averaged traces represent 2g (green, n=2 experiment, 6 cells) via FRET readout with the EPAC-S^H187^ sensor after 1 µM stimulation with Isoproterenol. Error bands represent sem.

**Figure S6:**
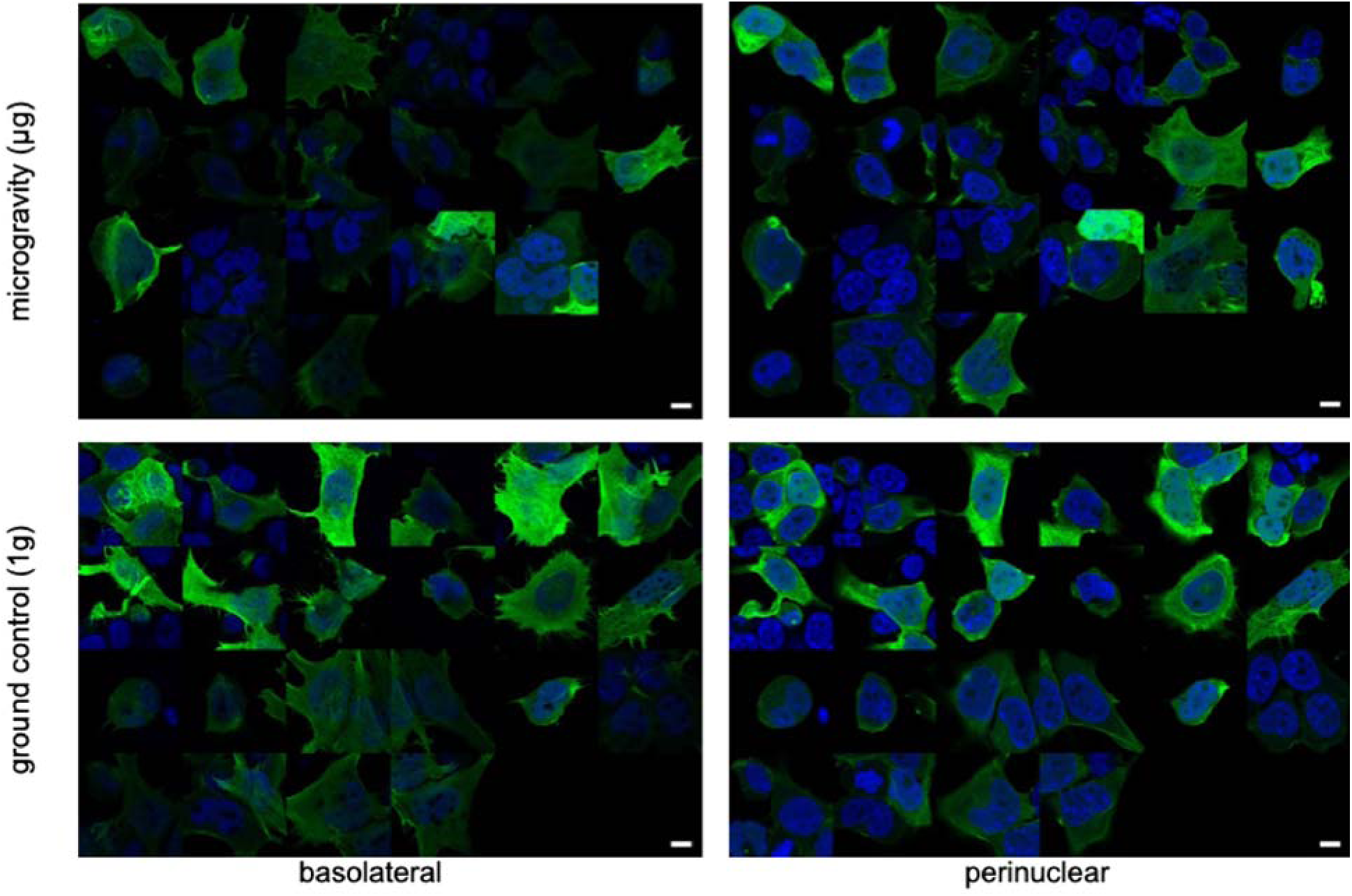
mosaic of HEK293 cells transfected with lifeact-EGFP and exposed to control gravity conditions or simulated microgravity for 24h. Representative images (24 per condition) from two experimental sessions conducted at the Ground Based Facility (GBF).

**Figure S7:**
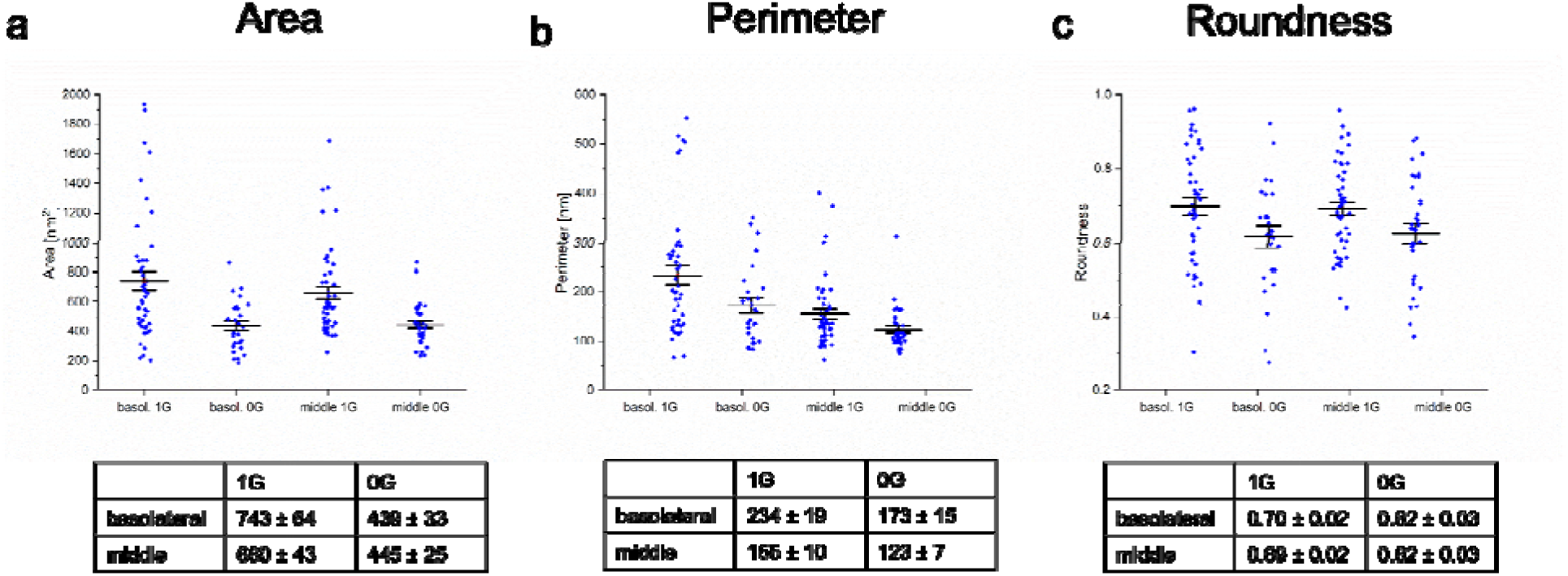
Qualitative morphological changes at HEK293 cells transfected with lifeact-EGFP and exposed to control gravity conditions or simulated microgravity for 24h. Displayed value indicate a) cell area within the confocal section, either at the basolateral membrane or at an intermediate perinuclear section; b) cell perimeter within the confocal section, either at the basolateral membrane or at an intermediate perinuclear section and c) overall cell roundness observed at the confocal section, either at the basolateral membrane or at an intermediate perinuclear section. Number of images acquired for basolateral µg: 27, 1g: 45; for middle µg: 34, 1g 48. The data originate from two experimental sessions conducted at the GBF.

**Figure S8:**
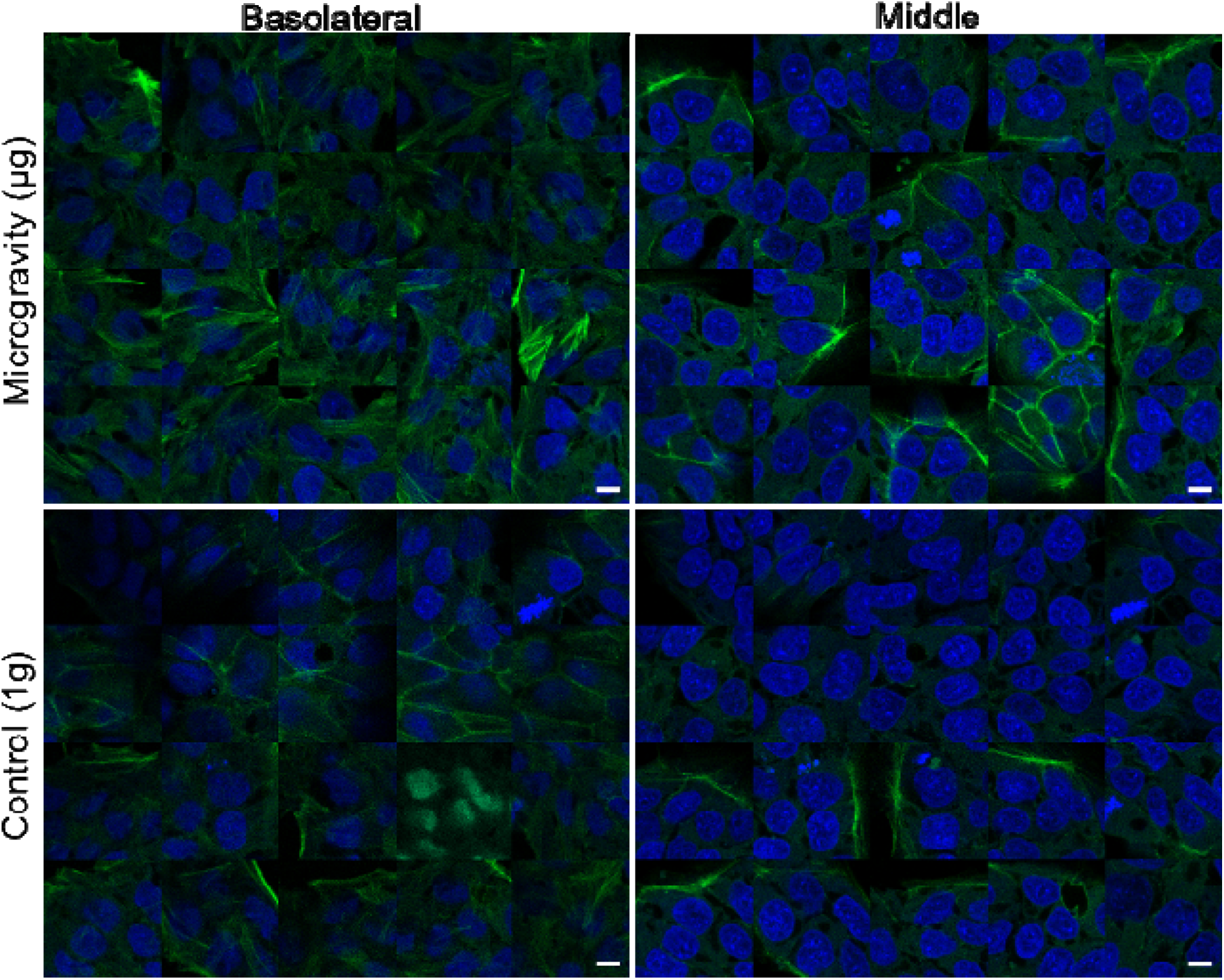
mosaic of hiPSCs cells expressing endogenously tagged EGFP-actin and exposed to control gravity conditions or simulated microgravity for 24h. Basolateral and perinuclear refer to the position where the images were taken in the confocal z-stack. Number of images acquired for each condition is 20. The data originate from one experimental session conducted at the GBF.

**Figure S9:**
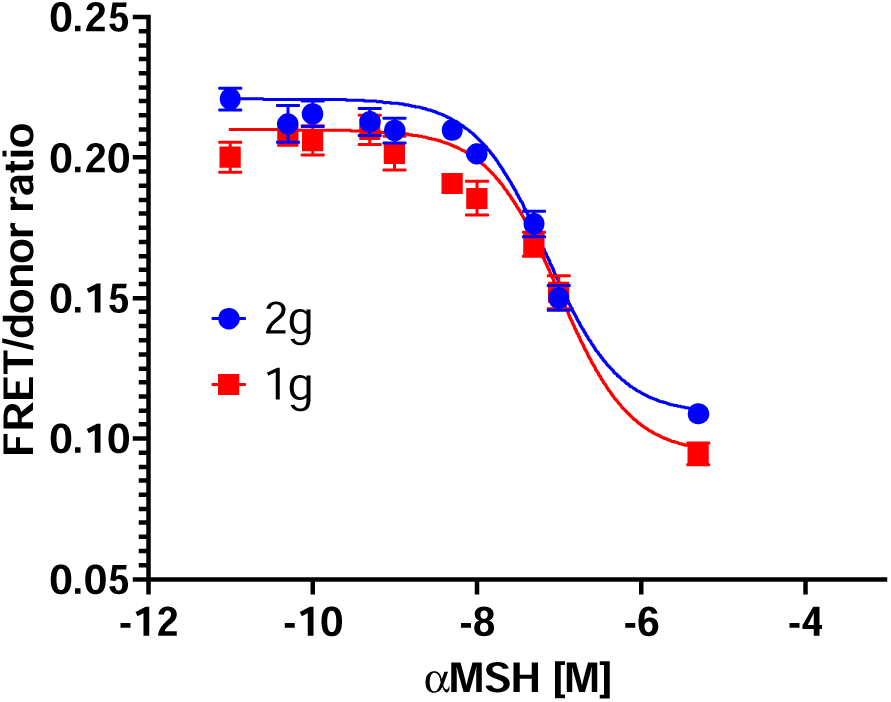
Effect of 24h hypergravity on cAMP production in HEK293 cells expressing the melanocortin 4 receptor (MC4R). The MC4R is a Gs-coupled a receptor involved in appetite regulation, upon stimulation with the agonist ⍰MSH. Here, concentration-response curve of FRET ratio to ⍰MSH stimulus does not highlight any significant difference between 2g and 1g control (1 biological replicate).

**Supplementary Table 1:** result of the fit of kinetic cAMP accumulation traces in Figures 2,4 and S5 and associated confidence intervals for the time constant τ.

| | $\tau$ [s] | 95% confidence interval from fit | | calculated standard error |
| --- | --- | --- | --- | --- |
|  |  | lower | upper |  |
| HEK 2g (Figure 2) | 9.671 | 8.869 | 10.47 | 0.41 |
| H9c2 2g (Figure 4) | 30.42 | 24.74 | 36.11 | 2.79 |
| H9c2 retinoic 2g (Figure S5) | 3.294 | 2.385 | 4.203 | 0.45 |
| HEK 1g (Figure 2) | 156.9 | 144.1 | 169.7 | 6.49 |
| H9c2 1g (Figure 4) | 21.44 | 20.01 | 22.88 | 0.72 |

